# Computational model of developmental self-assembly of a cortical microcircuit

**DOI:** 10.64898/2026.11.09.762492

**Authors:** Umar Abubacar, Cayla Harris, Ryan Bournes, Roman Bauer

## Abstract

Cortical microcircuits emerge from local developmental interactions, yet largescale computational models typically prescribe connectivity rather than explain its formation. Here we use BioDynaMo to simulate the self-assembly of ***∼***70,000 cortical neurons from 1,200 progenitors in 3D physical space through migration, neurite growth and contact-based synapse formation. With one calibrated filling fraction per connection type, the locally formed synapses reproduce experimentally measured connectivity across 64 connection types (mean absolute error 0.0062) without prescribing individual connections. The grown spiking networks sustain stable, inhibition-dominated activity, and the same developmental rules, given different targets, generate both a canonical cortical microcircuit and a visual-cortex variant. Inverting the migration programme produces a reeler-like column whose aberrant connectivity destabilises network activity. Together, these findings provide a mechanistic link between local developmental processes and the emergence of cortical anatomy, connectivity and function.

---

The mammalian genome specifies a cortex of extraordinary complexity, yet it is far too small to encode each connection individually [1]. Wiring must instead emerge from compact developmental rules unfolding in space and time, so the link from genotype to phenotype runs through development. Cortical neurons are born with layer-specific identities, migrate radially to their laminar positions, and extend axons and dendrites guided by molecular gradients. Synapses can form wherever these processes physically overlap, and cell-type-specific rules then determine which contacts become functional. These processes are stochastic, interacting and multiscale, so computational models are needed to test hypotheses about them.

Cortical microcircuits are the local networks through which the cortex processes and routes information [2]. Their recurrent excitation, inhibition and layer-specific connectivity shape sensory responses, modulate neuronal gain and timing [3], support plasticity [4], and determine how activity propagates between cortical regions [2]. Understanding how these circuits develop links genetic and cellular mechanisms to cortical computation, and may reveal how altered migration, neurite growth or synapse formation produces circuit dysfunction in neurodevelopmental and psychiatric disorders [5].

Here we choose to model the canonical cortical microcircuit, a stereotyped arrangement of excitatory and inhibitory populations across layers that recurs throughout neocortex [6]. Existing microcircuit models occupy two extremes. Point neuron models abstract morphology away and represent connectivity as a statistical matrix, integrated from anatomical and physiological measurements [7], so they cannot address how morphological properties constrain circuit function. Reconstruction-based models instead fix an adult snapshot, read from serial electron or light microscopy or prescribed from mapped cell-type densities [8–10], as do dense connectome reconstructions of retina and of invertebrate nervous systems [11–13]. All obtain connectivity independently of growth, either reading it from an imaged volume or prescribing it as a fitted probability, rather than from the developmental process that produced the morphology and the overlap constraint that process implies [14]. Neither reconstructions nor static models explain how this structure emerges. Developmental modelling can, by following the rapid rise in complexity from progenitors to a mature circuit.

Existing developmental models stop short of a microcircuit. Analytic and stochastic branching descriptions reproduce single-cell morphology in isolation [15]. Optimalwiring models grow arbours toward supplied carrier points and, grown competitively, neighbouring arbours tile the available space [16], but that substrate is prescribed rather than grown. Stochastic network generators extend this to populations and read connectivity off axo-dendritic proximity [17], but growth remains independent of neighbouring cells. Agent-based implementations address neurite growth under chemical guidance [18], gene-regulatory self-construction [19, 20] and Bayesian calibration against single-cell morphology [21], at the scale of single cells or small populations. To the best of our knowledge, no developmental model closes the loop against a measured population connectivity matrix at microcircuit scale.

Agent-based models are mechanistic. Each cell acts on local rules in physical space, interacting mechanically with its neighbours. We present an agent-based developmental model, built on the BioDynaMo simulation platform [22], in which *∼*70,000 cortical neurons self-assemble in three dimensions through gene-regulatory-network-controlled migration and growth-cone-guided neurite extension. Synapses form where grown axons and dendrites touch, and an optimiser identifies, for each connection type, the fraction of contacts that become synapses so that the grown neurons reproduce experimentally observed connectivity statistics. This contact-to-synapse step follows established practice [23, 24]. What is new is that the substrate itself is self-assembled, so the overlap statistics that bound the calibration are outputs of the growth rules rather than of experimental reconstructions.

Three things follow. First, the model is a multiscale developmental account of a canonical cortical microcircuit [6]. One simulation spans intracellular state, migration, neurite geometry, connectivity and spiking dynamics. Second, it shows how the complexity of cortical organisation can arise from simple rules [1]. Laminar architecture, morphological diversity and connectivity statistics are consequences of local developmental rules. Third, simple local rules are enough to produce a functioning microcircuit. The grown network matches the reference connectivity and supports stable spiking activity.

## 1 Results

### 1.1 Growth rules assemble a laminated cortical column

The column assembles through a sequence of developmental stages (Fig. 1a,b; Supplementary Video 1). Building on earlier agent-based approaches [18], no neuron is placed or wired by hand; layer depths and connection probabilities enter only as targets. A gene regulatory network first drives progenitor division through a sequence of progenitor states, so that laminar identity is fixed at birth order. The resulting neurons migrate radially along a simulated corticogenesis gradient. That migration reproduces the inside-out sequence of mammalian cortical development, in which deep layers form first and superficial layers last [20, 26]. Each neuron then extends axons and dendrites whose growth cones integrate local chemical gradients, directional momentum and stochastic exploration. The resulting layer-specific arbours constrain where synapses can physically form (Fig. 1c; Supplementary Video 2).

**Fig. 1.**
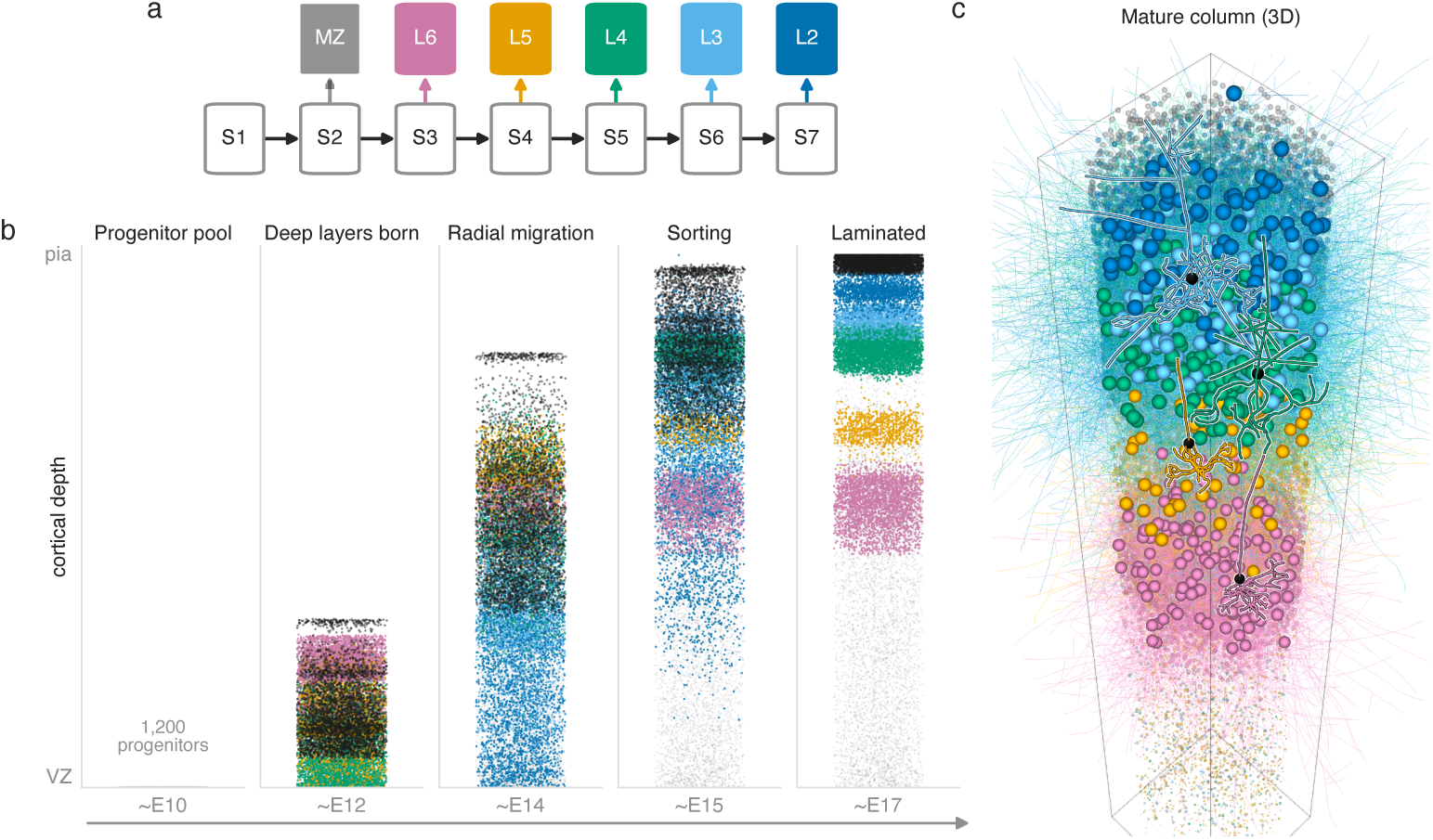
Cortical column self-assembly from developmental growth rules. (a) Gene-regulatory-network state machine. Progenitor states (S1–S7) gate layer-fate specification, assigning each neuron a laminar identity (marginal zone, L6–L2) at birth order, following Bauer et al. [20]. (b) Developmental time-lapse of the full-scale column over five stages (see also Supplementary Video 1). A progenitor pool proliferates, differentiates and migrates radially to produce the inside-out lamination observed in rodent neocortex, deep layers settling first [25]. Cells are coloured by GRN state, using the palette of panel a; dividing progenitors are mid grey, marginal-zone cells dark grey and cells that never become neurons light grey. (c) The mature column in three dimensions (perspective view), from a visualisation run of the same growth rules at reduced growth fraction, in which 508 neurons grow neurites. Somas (enlarged for visibility) and neurites are coloured by layer; neurites of one third of L2/3 cells are drawn, and segments above the pia are omitted. Highlighted: four excitatory neurons from the full-scale column (L3, L4, L5, L6; dendrites only), each the cell closest to its layer median across six morphometrics, placed at its layer’s median soma depth.

We implemented these stages on the BioDynaMo agent-based platform [22]. GRN-controlled division and differentiation turn 1,200 progenitors into a column of 70,328 neurons (84% excitatory, 16% inhibitory) bearing 280 million neurite segments, in 25 hours on a single high-memory compute node. Growth brings 182 billion pairs of processes close enough to touch, and the calibrated filling fractions convert a small minority of those into 205 million connected pairs for export to NEST.

Layers emerge in inside-out order, from the marginal zone at the pial surface to L6 at the white matter, with a soma span of 1,694 µm (Fig. 1b). Laminar depth targets are an input, taken per layer from rat barrel cortex measurements [27], and the migration parameters that meet them were tuned by hand rather than optimised. Five of the six layers settle within 100 µm of their target depth, the marginal zone within 2 µm, and L6 sits 153 µm superficial (Supplementary Table S1).

### 1.2 Grown morphologies match reference bands

We validate against two independent reference pools. The first is 482 Blue Brain Project reconstructions of rat somatosensory cortex [8]. The second is 243 NeuroMorpho.org reconstructions [28], 235 mouse and 8 cat, from 14 archives spanning occipital, somatosensory and frontal areas. We do not target any area, so this pool describes generic cortical morphology. Using two independent pools means no result rests on a single reference.

Our criterion is population-level plausibility, not a per-neuron match to any individual reconstruction (Fig. 2a). A metric passes when the simulated population median falls inside the reference [p5, p95] band. Across ten groups, layers 2 to 6 by excitatory and inhibitory class, the grown column passes 67 of 84 comparisons (79.8%) against the Blue Brain reference and 58 of 75 (77.3%) against NeuroMorpho (Fig. 2e). The core metrics are bifurcation count, total path length and maximum extent, shown for the basal dendrite in Fig. 2b–d.

**Fig. 2.**
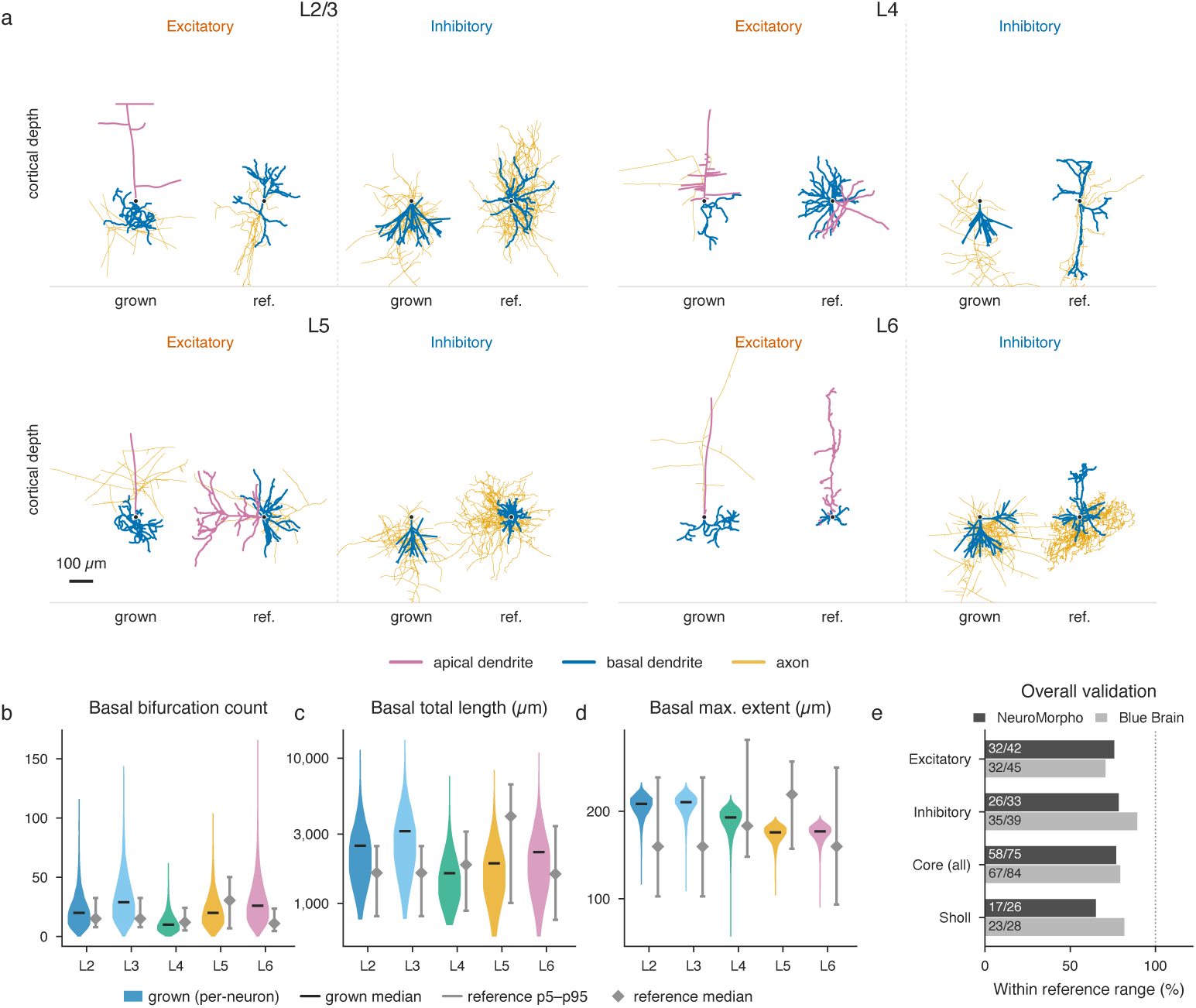
Morphology validation against the reference pool. (a) Representative grown and NeuroMorpho reference morphologies per layer (L2/3, L4, L5, L6), excitatory and inhibitory. Each grown cell is the closest available match to its reference partner across five morphometrics. Apical dendrite (pink), basal dendrite (blue), axon (orange). (b–d) Per-neuron basal-dendrite distributions, bifurcation count (b), total path length (c) and maximum extent (d), for all grown excitatory neurons of each layer (coloured violins) against the Blue Brain reference [*p*5*, p*95] range and median (grey). Basal dendrites are shown because they are present in every group of both references; apical and axonal results are summarised in e and in the text. (e) Pass rates for excitatory, inhibitory, all core metrics and Sholl profiles against each reference (NeuroMorpho, dark; Blue Brain, light). Reference groups are formed per layer, cell class and compartment across both pools, holding 3 to 77 reconstructions each (median 31).

Failures follow a depth gradient. Superficial excitatory axons are over-elaborated, branching and extending further than reconstructions, while deep-layer apical dendrites are under-elaborated, carrying less length and fewer branches. Layer-4 apical dendrites branch more than reconstructions. Six metrics fail against both references, mostly within this pattern. On 34 of the other 138 comparable tests, the grown neurons pass against one reference and fail against the other, because the references’ bands differ by up to 6.5*×*. Sparse deep arbours offer fewer cross-layer contacts, consistent with the connection types that reach the feasibility ceiling. Corrections that centred the apical medians collapsed their variance (layer-6 apical extent coefficient of variation 0.5%, against 43.5% in the reference), so we kept the more variable rules. Both references were scored on 500 grown neurons per layer and class, which gives the same core-metric verdicts as scoring all 69,859.

The simulation comprises two morphologically distinct inhibitory neuron types, perisomatic-targeting basket cells [29] and dendrite-targeting Martinotti cells with characteristic ascending, layer-1-projecting axons. Martinotti cells make up 26% of the column’s interneurons (2,894 of 11,300), after the somatostatin-expressing group of cortical interneurons [30]. The ascending Martinotti arbour later allows deep inhibitory-to-excitatory pathways to reach their targets. Deep interneuron axons follow the same depth gradient. Grown layer-6 interneuron axons branch less (5.8*×* fewer bifurcations than the reference median) and reach further (1.8*×*), a pattern weaker in layer 5 and absent in layer 4 (Fig. 2a). Inhibitory connection types are nonetheless the best recovered (Section 1.3). This grown morphology was then held fixed for all the results that follow.

### 1.3 One filling fraction per connection type fits the target

Growth completes before optimisation begins, so the axon–dendrite overlap between any two populations is fixed and each connection type’s probability depends only on its own filling fraction f*_ij_* and increases with it. The 64 cortico-cortical connection types therefore decouple into 64 independent one-dimensional fits.

We fit the 8*×*8 cortico-cortical filling-fraction matrix by gradient descent against the Potjans and Diesmann target matrix [7]. Per-iteration mean absolute error (MAE), the fitting objective over all 81 entries against the capped targets, fell from 0.0451 to 0.0040 over 55 iterations (91% reduction). Applying the best matrix at full scale gives per-connection-type MAE 0.0062 and RMSE 0.016. The Pearson correlation is r = 0.88 in log space over the 53 connection types that are non-zero in both matrices, and r = 0.98 linearly over all 64 (Fig. 3).

**Fig. 3.**
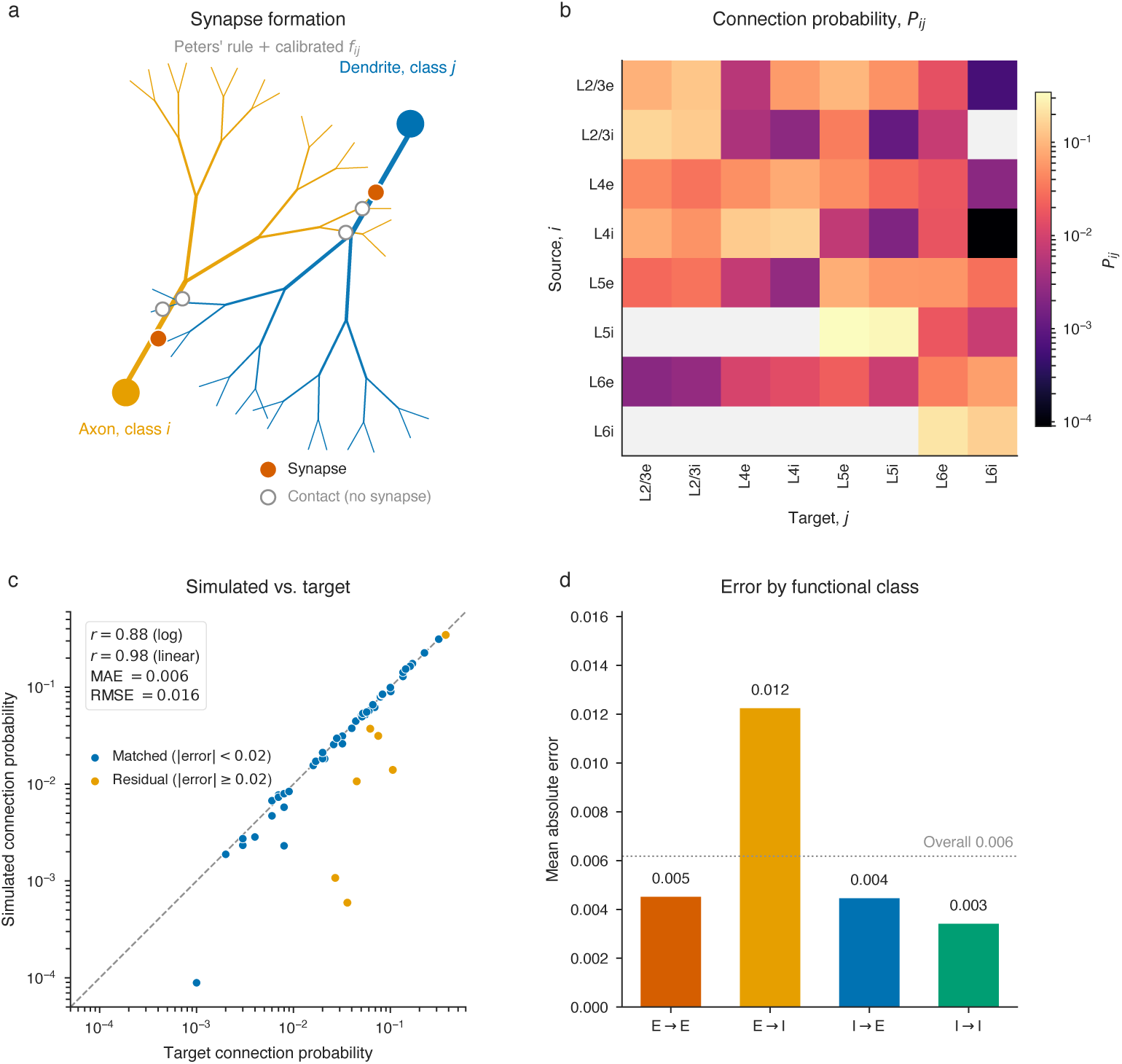
Connectivity calibration on the completed substrate. (a) Synapse-formation mechanism. Where an axon of class *i* overlaps a dendrite of class *j* (Peters’ rule), each contact becomes a functional synapse with the calibrated filling fraction *f_ij_*. (b) Calibrated connection-probability matrix *P_ij_* (source *×* target), on a logarithmic colour scale spanning 3 orders of magnitude. Grey cells carry no connection; all but one are types the target specifies as absent. The exception, L2/3I*→*L6I, has a non-zero target the substrate does not realise. (c) Simulated against target probability per connection type (log–log). Matched types (blue) lie within |error*| <* 0.02; larger residuals (orange) are mostly cross-layer feedforward types near the feasibility ceiling. Pearson *r* = 0.88 (log, 53 types nonzero in both) and 0.98 (linear, all 64); mean absolute error 0.0062, root-mean-square error 0.016. (d) Mean absolute error by functional class; dashed line, overall mean absolute error.

Mean absolute errors are 0.005 for E*→*E, 0.012 for E*→*I, 0.004 for I*→*E and 0.003 for I*→*I. The E*→*I class carries the largest error, concentrated in cross-layer feed-forward projections. The inhibitory classes are the best recovered, with local I*→*E connection types reaching 93–105% of target. The ascending Martinotti axon makes that possible by supplying overlap in layers the basket arbour does not reach.

Physical overlap bounds connectivity, so the grown morphology sets an upper bound on every connection type. Setting every filling fraction to one gives the largest probability each connection type can reach, a ceiling fixed by the morphology alone (Methods). The connection types that meet this ceiling are mostly long-range crosslayer excitatory projections, where the growth rules under-produce the axonal reach the target demands. L6E-to-L4I reaches 13% of its target and L5E-to-L2/3I reaches 42%. These are structural limits rather than tuning failures. No filling fraction recovers them, as their probability plateaus below target, and they cap the log-space correlation at r = 0.88 rather than at unity.

### 1.4 The same rules instantiate a second cortical area

Layer identity is assigned by the gene-regulatory network and arbour geometry by the growth-cone rules, and both are parameterised per layer rather than per area. To instantiate a second area we changed only the two inputs that are area-specific measurements, the connectivity matrix that the filling fractions are fitted to and the per-layer neuron fractions. The growth rules, the laminar depth targets and the morphology reference were not retuned for the visual area (Section 3.11). Any area-specific structure in the result therefore comes from those two inputs and not from retuned growth.

We tested this with a second variant, primary visual cortex. We used mouse V1 connection probabilities [31] as the target and left the growth rules at their canonical microcircuit values. The visual instance was grown at 25% of full scale, producing 18,115 neurons (15,924 excitatory, 2,191 inhibitory) and 3.8 million synapses. Morphological validation on 4,465 sampled neurons gives 55 of 75 core-metric passes (73.3%) against NeuroMorpho and 60 of 84 (71.4%) against Blue Brain. Its failures beyond those shared with the canonical microcircuit instance are mostly in axonal and layer-4 basal branching, whose parameters were tuned further after this instance was grown (Methods). The reference pool is not resolved by area, so this establishes that the visual morphologies stay within generic cortical bounds, not that they are visual-cortex morphologies.

We ran the same calibration on the visual substrate. Gradient descent converged in 64 iterations, from an MAE of 0.0251 to 0.0027 (Fig. 4a), reaching 0.0051 across the 64 cortico-cortical types shown in Fig. 4b, against 0.0062 for the canonical instance. The lower error reflects smaller target probabilities, averaging 0.032 against 0.053, since mean absolute error scales with the quantity being fitted.

**Fig. 4.**
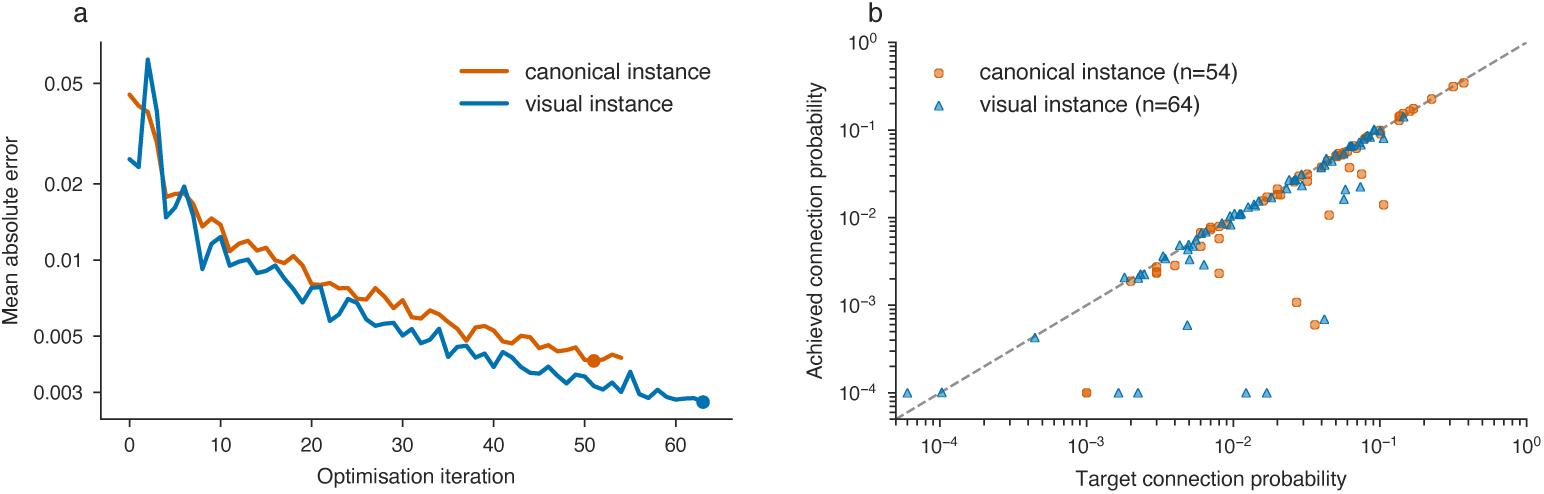
One set of growth rules, two cortical variants. (a) Filling-fraction optimisation for both instances, mean absolute error (fitting objective, against the capped targets) against iteration on a log scale. Filled markers mark the applied iteration. (b) Achieved against target connection probability for the cortico-cortical connection types of each instance, with the identity line dashed. Only connection types with a non-zero target are shown; achieved probabilities of zero are plotted at 10*^−^*^4^.

Both instances are constrained by the same morphology in the same way. The pathways that meet the feasibility ceiling fall in the same long-range cross-layer excitatory class in each, which is what a ceiling set by area-agnostic growth rules predicts. 13 of 64 visual connection types are structurally constrained.

We report the visual instance as a test of structural and calibration generality only. It was grown at reduced scale with a broad in-degree, both departures from the conditions the reference network’s downscaling assumes, so we do not report its dynamics.

### 1.5 Grown network sustains stable, inhibition-dominated activity

We simulated the calibrated canonical microcircuit instance in NEST [32]. We ran the published Potjans and Diesmann reference implementation size-matched to the grown column (Methods) and added the grown network as a second population wired from its own adjacency [7]. Both networks used identical single-neuron models, synaptic parameters and external background drive, so the networks differ only in recurrent connectivity and the delays derived from it (Fig. 5a,b).

**Fig. 5.**
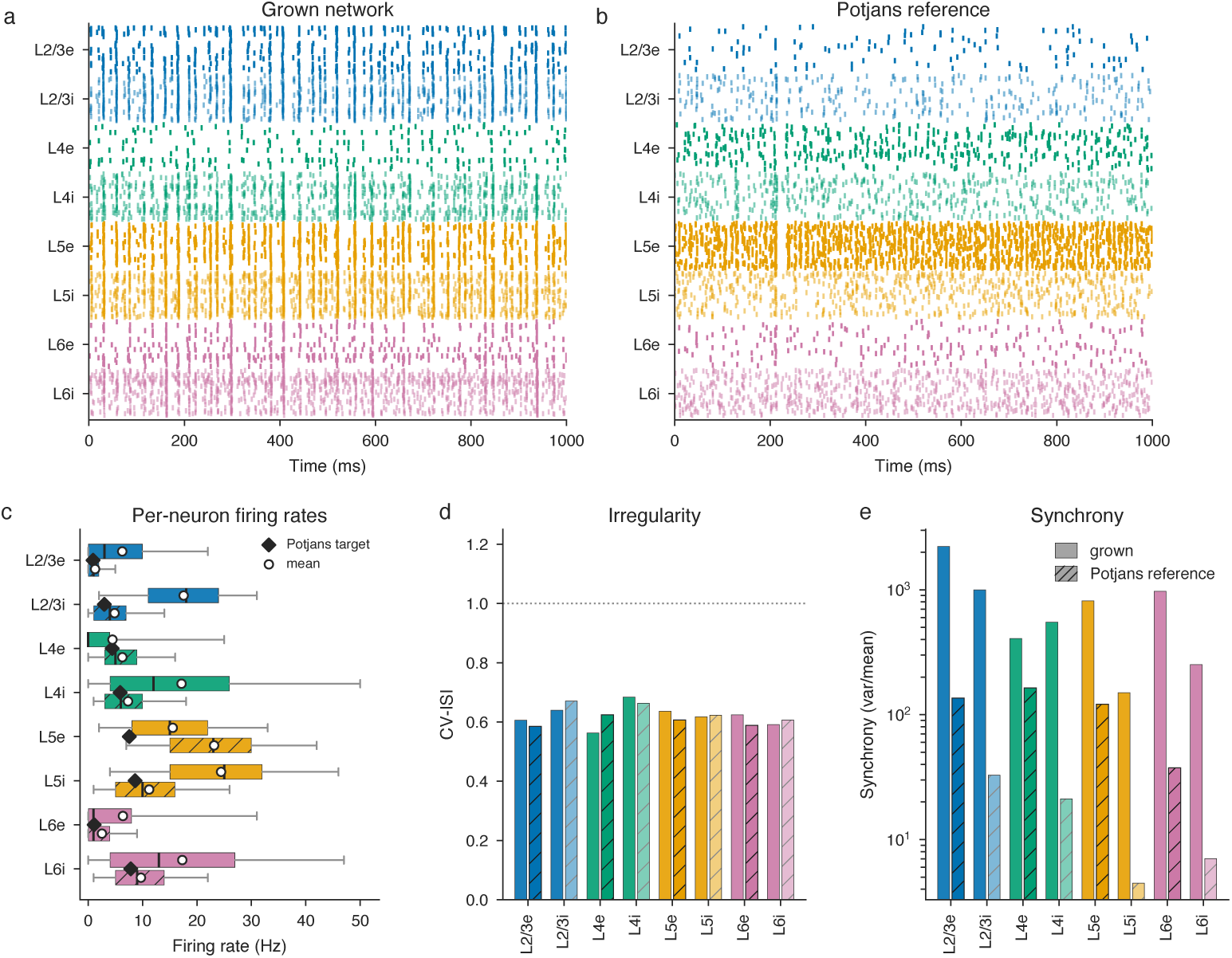
Spiking dynamics of the grown and the size-matched reference network. (a, b) Spike rasters of the grown and the size-matched reference network (random connectivity, matched *N* and *K* scaling, identical neuron parameters and external drive) over a 1,000 ms window. (c) Per-neuron firing rates per population (box, median and interquartile range; whiskers, 5th–95th percentile), pooled over 3 simulation seeds of the one grown connectivity. Layer 5 is the most active excitatory population in both networks, and inhibition exceeds excitation in every layer of the grown network. Both networks run above published rates under the same external-drive scaling, so absolute rates are not claimed. (d) Spike-train irregularity (CV-ISI); dotted line, Poisson; both networks are sub-Poisson. (e) Population synchrony (variance over mean of the 3 ms population spike-count histogram, log scale), higher in the grown network in every population over matched windows. Grown, solid; reference, hatched.

We measured population firing rates over n = 3 independent simulations of the grown network in a 1,000 ms analysis window (Fig. 5c, Methods). Layer 5 is the most active excitatory population in the grown network, in the size-matched reference and in the published profile. The remaining excitatory order differs, with layer 4 lowest in the grown network where the reference places it second. Inhibition exceeds excitation in all four layers of the grown network. The size-matched reference does not preserve this relation, with layer-5 excitatory firing above layer-5 inhibitory.

The grown network fires above the size-matched reference in six of the eight populations. The exceptions are layer-4 and layer-5 excitatory cells, at 4.47 and 15.55 Hz against 6.29 and 23.15 Hz. Grown connections follow morphological overlap, so the number of inputs per neuron varies widely, with a standard deviation of 27–67% of the mean, against 1.3–2.3% in the reference, where connections are drawn at a fixed probability. A few grown neurons therefore receive far more input than is typical.

Spike trains are irregular and sub-Poisson in both networks (CV-ISI 0.56–0.68 grown, 0.58–0.67 reference, Fig. 5d). Population synchrony is higher in the grown network in every population, by factors of 2.5 to 37.8 (Fig. 5e). Both networks were recorded over matched windows. The broad, right-skewed in-degree of the grown network may contribute, since neurons that share presynaptic input receive correlated drive.

### 1.6 Developmental and synaptic lesions give different phenotypes

Both stages that built the circuit, the developmental programme and the calibrated synapse-formation rule, remain available to perturb, and we trace the consequences of each to activity.

On the developmental axis, inverting the lamination programme regrows a reelerlike column [33] under identical growth and synapse-formation rules. This is a separate, single grown realisation, not the canonical column reported above. Mean connection probability rises from 0.049 to 0.078 (Fig. 6a–c, Supplementary Fig. S1). The rise is cross-layer, with cross-layer excitatory probability up 3.5*×* while within-layer types change little. Cross-layer excitatory connections join local inhibitory ones among the dominant types, and the column forms about 1.9*×* more synapses in total. The network loses stability, whereas the size-matched reference network at the same raised synapse count stays stable (3.4 to 33.4 Hz). This points to connectivity structure rather than synapse count as the cause, although it rests on a single grown realisation. The lesioned network settles at the refractory ceiling of the point-neuron substrate (500 Hz at τ_ref_ = 2 ms), so we report a loss of stability and do not quote firing rates for it.

**Fig. 6.**
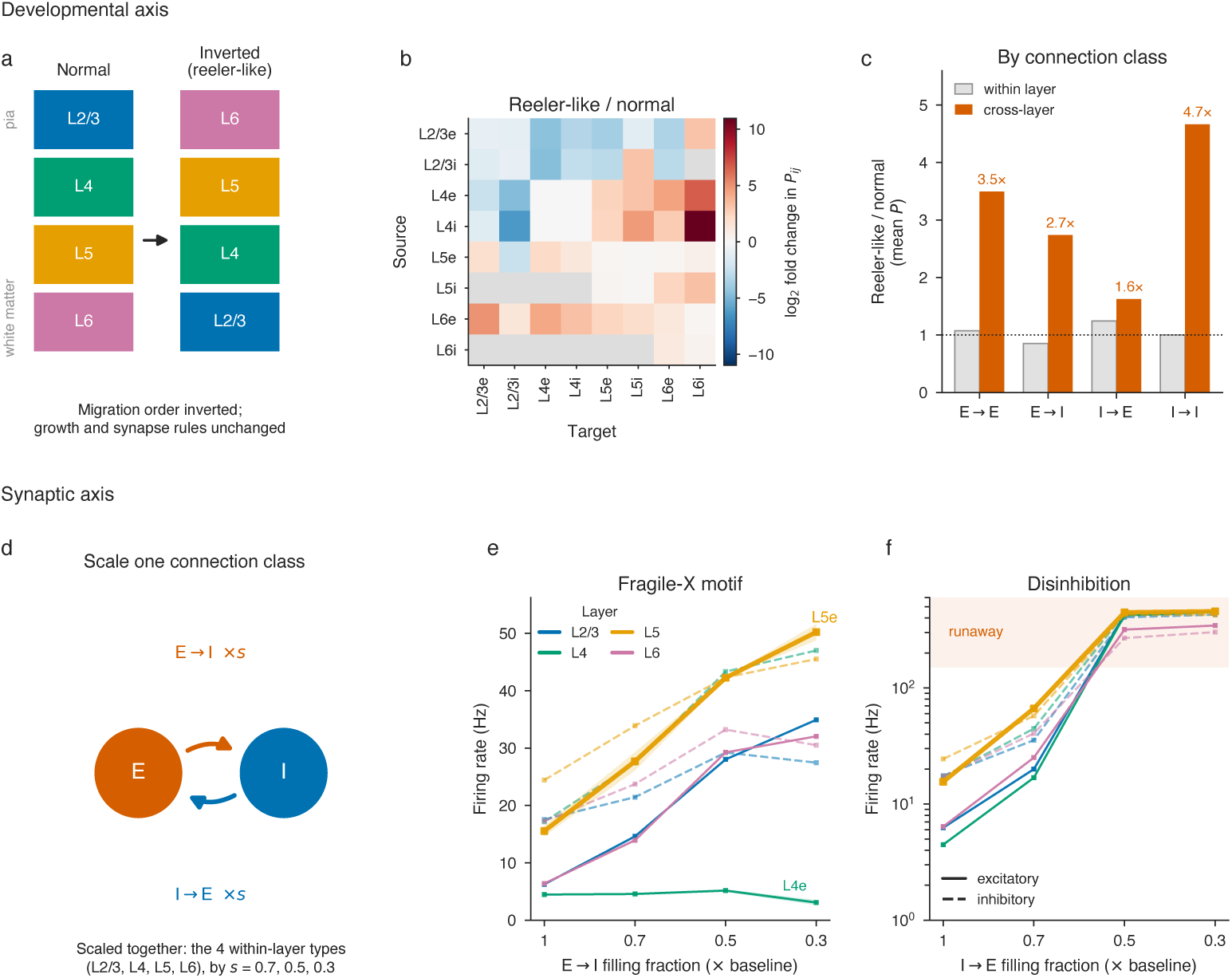
Developmental and synaptic perturbations of the grown column. (a–c) Developmental axis. (a) Inverting the migration order regrows a reeler-like column, with growth and synapse-formation rules unchanged (schematic). (b) Change in connection probability, reeler-like over normal (log_2_; grey, no connection in either). (c) Mean connection probability by class, reeler-like over normal, for within-layer (grey) and cross-layer (orange) connection types. The rise is concentrated in cross-layer types; cross-layer inhibitory types rise too, from much smaller baseline values. Single grown realisation of each column; full matrices in Supplementary Fig. S1. (d–f) Synaptic axis. (d) Perturbation scheme. The synapses of one class of connection, E*→*I or I*→*E, are thinned to a fraction *s* in the four within-layer connection types together, which alters connectivity and hence dynamics. Scaling E*→*I mimics the connection deficit reported in Fragile-X models; scaling I*→*E models disinhibition. (e) E*→*I knockdown (Fragile-X motif). Per-population firing rate against knockdown strength, showing graded hyperexcitability; colour shows layer. L4E is the only population that does not rise. (f) Disinhibition (I*→*E knockdown). Hypersensitive, with *×*0.7 graded and stronger knockdowns entering a runaway regime (shaded, log scale). Excitatory populations solid, inhibitory dashed. L5E and L4E emphasised. Points are the mean of 3 seeds; shading in (e) is *±*1 s.d. across seeds.

On the synaptic axis we thinned the synapses of individual connection types on the unchanged morphology and re-simulated in NEST (Fig. 6d–f). Weakening excitatory drive onto inhibitory interneurons produced a graded hyperexcitability. This is an excitation/inhibition-imbalance motif of Fragile-X syndrome, an inherited neurodevelopmental disorder caused by loss of the FMR1 gene product [34, 35]. Layer-5 excitatory firing rose steadily with knockdown strength, from 15.55 Hz at baseline to 50.2 Hz at *×*0.3 (Fig. 6e). Layer-4 excitatory cells are the only population that does not rise, ending at 3.1 Hz against a baseline of 4.47 Hz while every other population increases by 1.6 to 5.6*×*. The knockdown is applied to the four within-layer connection types only. The *×*0.5 level halves these connections, matching the roughly halved excitatory drive onto fast-spiking interneurons in layer 4 of mice lacking the gene (Fmr1 knockout), where connections are weaker and, at two weeks, fewer [34]; we apply it within every layer. Applied to all cross-layer pairs, it saturates the network instead. The model thus matches the knockout’s hyperexcitability in direction but not in layer 4, the layer those recordings come from, where evoked activity in excitatory cells lasts longer [34]. It also omits the knockout’s more excitable excitatory cells.

Weakening inhibitory drive onto excitatory cells, raising the excitation/inhibition ratio [5], was instead hypersensitive. A *×*0.7 reduction already drove layer-5 excitatory firing to 66.5 Hz, and stronger knockdowns drove firing to about 456 Hz, near the model’s refractory ceiling, which marks a loss of stability rather than a meaningful firing rate. We draw the disease comparison from the graded regime and not from the runaway one. Each condition is the mean of 3 independent seeds, which agree to within 15% of the mean on every population.

## 2 Discussion

We present a developmental model in which cortical microcircuit connectivity emerges rather than being prescribed. Neuronal identities arise through gene-regulatory fate specification [20], neurites extend along spatial guidance cues, and synapses form where compatible processes meet [19], calibrated by one interpretable filling fraction per connection type. Each stage stays accessible after development, so a perturbation of any one can be traced through morphology, connectivity and activity.

Existing large-scale cortical models begin with prescribed connectivity, derived by touch detection and statistical pruning on reconstructed morphologies [8, 24], by distance-dependent rules [31], or by reconstructing the circuit of one animal [9]. They are more accurate cell by cell but capture only one circuit state. The present model is a framework for asking how that structure emerges and how perturbing fate specification, lamination, neurite geometry or synapse survival propagates into connectivity and activity.

Because the substrate is grown, it can be perturbed on two axes. On the developmental axis, inverting the lamination programme regrows a reeler-like column under unchanged growth and synapse-formation rules, and the aberrant cross-layer excitatory connectivity that results destabilises the circuit. On the synaptic axis, weakening excitatory-to-inhibitory connections produces a graded hyperexcitability, the direction reported in Fragile-X models, and weakening the inverse connections a far more sensitive disinhibition [5, 35]. Neither synaptic manipulation required retuning of single-neuron or synaptic parameters.

Neurodevelopmental disorders arise from altered development, and agent-based models have shown how altered developmental mechanisms recapitulate pathological properties of polymicrogyria [20]. Simulating each stage of circuit formation can help identify whether a phenotype originates in fate specification, migration, neurite growth or synapse formation, and trace how it reaches mature connectivity and activity.

Most morphological metrics fall inside the reference range, and the main discrepancies are shorter deep-layer apical dendrites and over-branched superficial axons. The growth rules are phenomenological, their parameters can be calibrated further (Methods), and future versions could constrain them with omics data [36].

The clearest target for the next version of the growth rules is axonal reach. Long-range cross-layer excitatory connections need more reach than the current rules produce, so those connection types stay below target at any filling fraction (13 of 64 in the visual instance), and the feasibility analysis identifies them before any fitting.

The spiking model uses point neurons and so ignores the grown dendritic detail. Each connection is also a single synapse rather than the multiple contacts of a biological one. Calibration matches connection probability per type, not each neuron’s in-degree. Constraining the in-degree distribution, or adding activity-dependent pruning, is a natural next step.

No input from outside the column is modelled, whether from the thalamus, basal ganglia or other subcortical structures, or from interareal, callosal and long-range horizontal connections. The cell-type repertoire is reduced to two inhibitory morphological classes and one excitatory morphology per layer. We report a single grown realisation of each column (canonical, visual and reeler-like), so across-realisation variability is unquantified.

A fully developmental model opens questions that models beginning with an adult circuit cannot address. Activity-dependent refinement, driven by spontaneous activity and early sensory input, would let experience shape the grown wiring and its mature function, extending earlier small-scale models that combined growth with homeo-static and Hebbian plasticity [19]. Because the stages through which genes shape cortex are explicit, gene-expression data could inform the rules for cell fate, migration, neurite growth and synapse formation, giving a mechanistic path from genotype to circuit phenotype. Disorder-associated phenotypes could then emerge from perturbed development instead of being imposed on a mature network. BioDynaMo scales to a billion agents on one server [22], so the same rules could extend toward a cortical area, interacting areas and ultimately a whole brain, with its folding pattern [37] and spontaneous activity such as retinal waves [19]. Because the complexity arises from simple local rules, it need not be specified in advance, setting the approach apart from reconstruction-based models [8]. Across areas and species, they will trace differences between circuits to differences in how they develop, elucidating how the genome encodes developmental rules rather than wiring [1].

## 3 Methods

### 3.1 Simulation framework

All simulations used the BioDynaMo agent-based framework [22], which provides efficient parallel execution of biological growth models. Following the formulation introduced by CX3D [18], each cell carries a small set of biological primitives. It can divide, move, extend and branch neurites, and detect local chemical substances. The developmental sequence emerges from these primitives, with no top-down specification of layer positions or wiring.

The simulated column is a cylinder of radius approximately 250 µm spanning the full pial-to-white-matter depth of the domain. The Y axis represents cortical depth. Neurons migrate within this radial constraint, which relaxes once lamination completes, so horizontal extent varies between stochastic runs. We ran the full-scale simulation for 15,000 timesteps at a growth fraction of 0.3, the parameter that sets what proportion of the layer-committed cells become neurons (Section 3.2). The run that produced the reported network completed in 25 hours on a single node using 64 cores and a measured peak of approximately 210 GB of memory.

Three developmental stages mirror biological corticogenesis. Migration sets neuronal position from a per-type migration substance, extending to every layer the threshold rule that Bauer et al. apply to marginal-zone cells [20]. Each neuron is assigned a substance level by its layer identity and migrates until that level is reached, so laminar position follows from fate rather than from a prescribed distance. Outgrowth extends axons, basal dendrites and apical dendrites under compartment-specific growth rules. Synapse formation converts geometric axon-to-dendrite contacts into synapses under Peters’ rule [14], as detailed below. Morphologies are stored in standard SWC format.

### 3.2 Gene regulatory network and neurogenesis

Cell fate follows the gene regulatory network (GRN) model of Bauer et al. [20], in which a sequence of progenitor states, advanced by an internal clock, controls birth order. Progenitors in the ventricular zone first undergo symmetric proliferative divisions that expand the pool. They then switch to asymmetric neurogenic divisions producing one progenitor and one post-mitotic neuron. Each neuron inherits the layer identity encoded by its mother progenitor’s GRN state at division. Early-born neurons therefore take deep-layer identities (L6, L5), and progressively later-born neurons more superficial ones (L4, L3, L2), mirroring the temporal cascade of mammalian corticogenesis [26].

Starting from 1,200 progenitors, the programme expands the population to 258,950 committed cells. Marginal-zone cells aside, the gene regulatory network assigns each a layer identity. The growth fraction then sets how many of them become neurons, split between an excitatory and an inhibitory path. At the value used here 28.5% do so, yielding the 70,328-neuron network (59,028 excitatory, 11,300 inhibitory). The remainder stand in for cells that undergo apoptosis, adopt glial fates or become cell types not modelled here. We calibrated the growth fraction once against the population sizes of the Potjans and Diesmann reference column [7].

### 3.3 Guidance substance

A single static gradient guides neurite growth, defined on BioDynaMo’s Cartesian concentration grid [22]. The guidance field is a Gaussian band centred on the pial surface (σ = 640 µm), so only its lower half lies within the column, initialised once at the start of the simulation and neither diffusing nor decaying thereafter. It gives a smooth axial profile, high at the pia and low at the white matter. It stands for a pia-to-white-matter morphogen gradient and is not identified with a specific molecule. The axonal guidance term points towards white matter and the apical term towards the pia; whether an axon descends or ascends is set by its target (below). The gradient weight w*_grad_*, specified per cell type and layer, sets how strongly each compartment is biased relative to its momentum and stochastic terms.

Axon termination uses a geometric target rather than a second chemical cue. Each axon is assigned a target depth and growth direction at construction, sampled per cell type. The ascending fraction is 0% in L2, 30% in L3, 80% in L4, 40% in L5 and 30% in L6, and every Martinotti axon ascends. While extending, it obeys the same three terms as any other neurite. Once the tip crosses its target depth, bifurcation and diameter-loss rates rise several-fold, forming a compact terminal arbour at that depth. In dendrite-targeting Martinotti axons this phase also spreads horizontally, giving that class its layer-1 tuft. One rule therefore produces both the long vertical projections and the terminal arbours of cortical axons.

### 3.4 Neurite growth models

Each neurite tip is modelled as a growth cone, a motile sensory structure that samples its local chemical environment and steers extension accordingly. At each timestep the growth cone integrates three competing influences into a new extension direction,

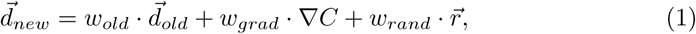

where 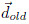 is the previous direction, giving momentum analogous to cytoskeletal persistence. The term *∇*C is the local gradient sampled at the tip, and 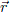 is a random unit vector giving stochastic exploration. The weights w*_old_*, w*_grad_* and w*_rand_* are unnormalised and specified per layer and per cell type, which is what lets compartment-specific growth patterns emerge from one mechanism.

The tip then advances by a compartment-specific step length of 10 µm to 40 µm, and diameter decreases by a small attenuation factor. Growth terminates when diameter falls below a compartment-specific threshold, naturally producing the tapering of biological axons and dendrites. This approach follows Duswald et al. [21], extended here with compartment-specific branching rules and a tufting mechanism for apical dendrites.

Direction weights, per-layer bifurcation probabilities and per-layer diameter-loss rates constitute the full morphological parameter set and are tabulated in Supplementary Tables S2 and S3. Weights were held fixed across cell types except where a distinct value was needed for a compartment– or subtype-specific morphology, so basal, apical, basket and Martinotti compartments each carry their own momentum and random weights. Bifurcation probability and diameter-loss rate are the principal levers tuned during morphology fitting.

#### 3.4.1 Axon growth

Axons grow with high directional momentum (w*_old_* = 12.5 for excitatory neurons), a layer-specific gradient weight (w*_grad_* = 1.2 to 2.0) and a small stochastic term (w*_rand_* = 0.8). The guidance term points towards white matter; an axon assigned an ascending target climbs on its initial direction and high momentum, so the same weights produce both descending and ascending projections. Branching occurs with a per-layer probability per timestep, creating asymmetric side branches while the main trunk continues. A new branch point requires a minimum distance from the previous one (70 µm for excitatory, 20 µm for basket and 40 µm for Martinotti axons). The probability is scaled by 0.6 on trunks and 0.4 on excitatory side branches, and once an axon passes its target depth, branching and diameter loss rise fivefold (twofold for Martinotti cells) to form a terminal arbour. Diameter attenuation naturally yields a proximal-dense branching pattern, because thicker initial segments support more branching events before reaching the termination threshold (0.3 µm). Daughter diameter is reduced by an empirical taper factor of 0.8 (0.85, with termination at 0.15 µm, in layer-6 interneurons). Bifurcation probability, diameter attenuation and direction weights are parameterised independently per layer.

#### 3.4.2 Apical dendrite growth

Apical dendrites grow towards the pia along the positive guidance gradient with high persistence (w*_old_* = 8.0), weak gradient coupling (w*_grad_* = 0.3) and low randomness (w*_rand_* = 0.3). The result is the straight trunk characteristic of pyramidal cells. When guidance-substance concentration exceeds a threshold in L1 and the marginal zone, a tufting zone activates. Bifurcation probability rises 100– to 200-fold depending on layer, and branching switches from asymmetric lateral twigs to symmetric horizontal bifurcation, producing the terminal tuft. Pre-tufting oblique branches undergo accelerated diameter loss, at 5*×* the trunk rate, so they terminate as short lateral twigs. Daughter diameters follow Rall’s law, whose symmetric-bifurcation form gives a factor of 0.63.

#### 3.4.3 Basal dendrite growth

Basal dendrites grow with minimal gradient influence (w*_grad_* = 0.03), reflecting their local arborisation. High momentum (w*_old_* = 6.0) keeps segment trajectories straight between branch points, while a comparatively large stochastic weight (w*_rand_* = 2.0) and symmetric bifurcation produce the radiating arbour. All parameters are per layer and were tuned against the reference pool.

#### 3.4.4 Inhibitory neuron growth

Inhibitory neurons, 16% of the grown population, use the same rules with a distinct parameterisation. Each grows five basal dendrites in fixed radially symmetric orientations and a single axon. Basket cells grow no apical dendrite, whereas every Martinotti cell grows one dendrite directed towards the upper layers, approximating the ascending tuft of their usually bitufted dendrites [29]. We model two classes, perisomatic-targeting basket cells and dendrite-targeting Martinotti cells [38], capturing the coarse perisomatic versus dendritic distinction. Finer interneuron subtypes are not represented.

Inhibitory axons have lower momentum (w*_old_* = 5.5 against 12.5 for excitatory) and higher stochasticity (w*_rand_* = 1.5), producing local bushy arbours rather than long-range projections. Higher diameter-loss rates keep them spatially compact, consistent with the local connectivity of cortical interneurons [30]. Martinotti axons carry their own weights (w*_old_* = 8.0, w*_rand_* = 1.5) together with the ascending target depth that produces the layer-1 tuft. We set the basket-to-Martinotti ratio so Martinotti cells comprise 26% of interneurons (2,894 of 11,300 across L2 to L6). That approximates the somatostatin-expressing group, one of the three non-overlapping molecular classes that together account for nearly all neocortical interneurons [30]. Because the model represents only two inhibitory classes, the remaining 74% are grown as basket cells, which absorbs the parvalbumin-expressing basket population together with the 5HT3aR-expressing group that is not separately represented.

### 3.5 Reference data sources

We use two morphology reference pools. The first is 482 reconstructions from the Blue Brain Project somatosensory dataset [8], of which 303 carry both a layer and a cell-class label and enter the comparison. This pool is area-specific, and it gives the deepest coverage where the grown column is hardest to validate, at 59 layer-5 and 77 layer-6 excitatory cells.

The second is 243 cortical reconstructions from the NeuroMorpho.org archive [28], with per-neuron provenance provided in the accompanying dataset. It comprises 235 mouse and 8 cat neurons from 14 contributing archives. No reconstruction was removed from the pool on the basis of validation results. The largest single source is the Allen Cell Types database [39], with 90 neurons. Regions span occipital (121), somatosensory (67) and frontal or prefrontal (54) cortex, with 1 presubicular neuron. Ages are predominantly adult (172) or young adult (50), with 11 young and 10 not reported.

Pooling across areas and archives is deliberate. We do not target any area, so this pool describes generic cortical morphology rather than the morphology of one region, and the same rules can grow more than one instance.

Layer and cell class are taken from each pool’s own per-neuron record. For NeuroMorpho this is each reconstruction’s NeuroMorpho.org entry, and cells recorded as layer 2/3 count in both the L2 and L3 groups. Groups are formed per layer, cell class and compartment, and a group holding fewer than three reconstructions is excluded rather than scored. Both pools are thinnest in the inhibitory apical compartment.

The thalamus is not modelled, so the 17 thalamic pathways are excluded from the fitted error. Connectivity targets for the canonical microcircuit instance come from Potjans and Diesmann [7], which compiled cell-type-specific connection probabilities from combined cat and rat data. Targets for the visual instance come from the Allen Institute mouse primary visual cortex model [31]. That model reports peak connection probabilities with a Gaussian dependence on lateral distance whose width differs by pair type. We averaged each profile over the distribution of distances between random points in a disk of the column radius (250 µm), collapsed the seventeen Allen cell classes onto the eight modelled ones and excluded L1, giving the 64 cortico-cortical targets. The grown L2 and L3 share the single L2/3 target. The resulting values are column averages and are therefore smaller than the peak probabilities the model reports. In both cases the target matrix is an input to calibration and is never used during growth.

### 3.6 Morphometric analysis

We separated morphologies into axon, basal dendrite and apical dendrite compartments. For each compartment we computed bifurcation count, total path length and maximum Euclidean distance from the soma. These three scalars are the core metrics and carry the primary pass or fail criteria.

Sholl analysis [40] quantifies radial complexity by counting intersections with concentric spheres centred on the soma, at radii from 10 µm to 400 µm in 10 µm steps. Unlike the core metrics, which use medians, Sholl profiles are compared using population means, which preserves curve shape. Normalising each curve by its maximum removes absolute scale dependence and allows shape comparison between neurons of different sizes. That normalisation is necessary because the simulated population spans a wider range of total arbour length than the reference pool.

As supplementary diagnostics we compute the spatial envelope of each compartment. This comprises horizontal extent (maximum distance from soma in the XZ plane), vertical extent (maximum distance along Y) and the horizontal-to-vertical (H/V) ratio. The H/V ratio separates laterally spreading arbours (H/V > 1) from vertically oriented ones (H/V < 1). These are reported but are not pass or fail criteria.

### 3.7 Validation methodology

Unlike formal Bayesian calibration, which infers parameter posteriors for a single growth model against one morphological class [21], we tune parameters for all cell types and compartments of a full column iteratively, without formal inference. Each round runs a simulation, compares morphometric statistics to the reference pool and closes the gaps that comparison identifies, until the statistics converge. The remaining discrepancies, chiefly short deep-layer apical dendrites and over-branched superficial axons, lie in parameter values rather than in the form of the growth rules. Each compartment’s branching, extension and attenuation are free parameters, so formal Bayesian calibration per compartment [21] is a direct route to reducing them, and one of the next steps for this model. For the deep apical dendrites it will need to preserve the variability the reference shows.

Reference groups are too small to support a meaningful held-out split, so we report pass rates on the full reference set. The reported pass rates therefore describe the fit between a 70,328-neuron simulation median and a reference distribution of at most a few tens of neurons. They are not test-set generalisation. Generalisation to unseen morphology datasets is left to future work.

The primary criterion requires the simulated population median to fall within the reference [p_5_, p_95_] percentile band. The core-metric test set spans three metrics, three compartments and ten cell types, giving 90 pooled comparisons. We drop tests whose reference group holds fewer than three reconstructions, leaving 75 for NeuroMorpho and 84 for Blue Brain. As a stricter alternative we also report how many comparisons have a simulated-to-reference median ratio inside the [0.7, 1.5] median ratio, which holds for 37 of 75 NeuroMorpho and 42 of 84 Blue Brain comparisons.

A Sholl curve passes if at least 70% of active radii fall within a band of the reference mean plus or minus 1.5 standard deviations. That band covers roughly 87% of a normal range and is deliberately wide, so as to absorb the variability of a small reference sample. We preferred it to a Wasserstein distance because it accounts explicitly for that variability, which in this pool often approaches the reference mean in magnitude.

Formal distributional tests are the wrong instrument at this sample asymmetry, which is *∼*70,000 simulated neurons against reference groups of 3 to 77 neurons. With that many simulated cells a difference far smaller than the biological variability registers as significant, so such a test measures statistical power rather than biological mismatch. We therefore validate on the percentile band supported by effect sizes, not on distributional identity.

### 3.8 Known artefacts of the reference pool

Three systematic artefacts bound what the morphology comparison can establish. All are properties of the reference pool or the domain, not of the growth rules.

The first is slice truncation. The pool draws on 14 archives whose slice protocols and axonal completeness differ and are not recorded per neuron. The simulation instead grows a full column, with a soma span of 1,694 µm. Any reconstruction prepared from tissue slices truncates arbours wider than the slice. Large basket-cell axons are that case: they reach neighbouring and distant columns and inhibit neurons in upper and lower layers [29]. Full-column arbours are therefore expected to exceed the reference pool at the widest radii, where the axon maximum-extent and outer-radius Sholl comparisons sit. Normalised measures such as Sholl shape and bifurcation ratios are less sensitive to truncation, and we treat them as the more reliable targets. We claim no quantified truncation correction, because the pool does not support one.

The second is column width. Basket-cell H/V ratios in the grown population are 0.71 to 1.02, indicating near-isotropic local arbours. Large basket cells instead spread their axons across neighbouring and distant columns, and are the main source of lateral inhibition within their own layer [29]. The cylinder caps horizontal extent near 250 µm, so recovering that anisotropy would need a tissue volume wider than one column.

The third is inconsistent compartment labelling. Neurite type is recorded per archive, and some reconstructions label no apical compartment at all, even for pyramidal cells that plainly have one. The layer-2/3 excitatory reference in Fig. 2a is one such case, which is why no apical dendrite is drawn for it. Group statistics are computed per compartment, so an unlabelled apical removes that neuron from the apical comparison rather than entering it as a zero. Single-neuron panels are the only place the omission is visible.

### 3.9 Synapse formation and filling fraction optimisation

Contacts are identified geometrically wherever an axonal and a dendritic segment from different neurons lie within 10 µm, implementing Peters’ rule [14]. Each contact is then independently converted to a functional synapse with probability f*_ij_*, where i and j index source and target population. In principle this can place several synapses on a pair whose arbours overlap at many sites. The reported network caps each connected pair at 1 synapse, a simplification consistent with fitting connection probability. Independent per-contact conversion of this kind, fitted to connection probability, does not also reproduce the number of synapses per connection, and Reimann et al. add a multi-synapse pruning stage precisely to constrain it [24]. We do not attempt it, because the point-neuron simulation represents each connection by a single synapse, so the number of synapses per connection never enters it.

Peters’ rule alone cannot account for the cell-type specificity of cortical connectivity, because neurons do not form synapses at every geometrically available contact. The realised share, the *synaptic filling fraction* [23], is well below unity and varies between connection types [24], reflecting molecular recognition, activity-dependent pruning and developmental stabilisation.

The filling fraction f*_ij_ ∈* [0, 1] is therefore a per-population-pair parameter rather than a global constant. Arranged across the eight cortical populations of layers 2/3, 4, 5 and 6 plus the thalamic population of the Potjans and Diesmann layout, these values form the 9*×*9 filling-fraction matrix, whose 8*×*8 cortico-cortical sub-block holds the 64 fitted connection types. The grown column contains no thalamic neurons, so the thalamic row and column are not fitted and sit at a negligible floor of 5 *×* 10*^−^*^5^. A pair sharing k contacts connects with probability

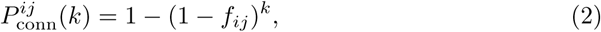

the exact per-pair probability.

The connection probability of a type is the average of this per-pair probability over all of its neuron pairs,

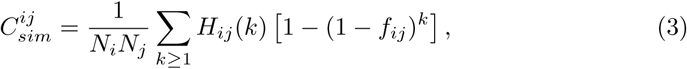

where N*_i_* and N*_j_* are the population sizes and H*_ij_*(k) counts the neuron pairs of that type sharing exactly k contacts. The histograms are counted once on the completed substrate, so each fitting iteration evaluates this sum in about 0.2 s instead of resampling synapses. When k f*_ij_ ≪* 1 it reduces to 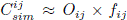, where O*_ij_* is the mean number of contacts per neuron pair, fixed by the morphological substrate. This separates the structural constraint O*_ij_*, set by morphology, from the molecular specificity f*_ij_*, set by optimisation. Because many neuron pairs share several contacts, the fit uses the full average rather than this approximation. The optimised f*_ij_* are functionally derived filling fractions, obtained from connectivity targets rather than from direct anatomical measurement [23].

Stochastic growth is strictly completed before optimisation begins. Once growth finishes the substrate no longer changes, which makes O*_ij_* deterministic, and gradient descent then adjusts the filling fractions against that fixed substrate. No morphological parameter is altered at this stage.

The loss is the sum of squared errors between simulated and target connection probabilities,

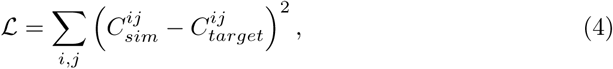

and its gradient with respect to the filling fractions is approximated as

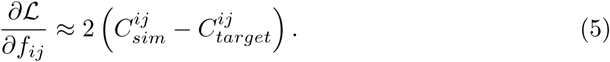

This residual form suffices on a static substrate, because there each connection type’s connectivity is monotone in its own filling fraction alone. The residual therefore points in the direction that reduces the error, and Adam scales it by its adaptive step rather than by the exact slope.

The Adam optimiser [41] provides the update

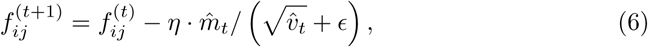

with learning rate η = 0.01 and bounds f*_ij_ ∈* [0, 1] enforced by clipping after each update. We preferred Adam to proportional rescaling because its momentum and adaptive step size prevent oscillation near the feasibility ceiling and give a well-defined convergence criterion. Optimisation stopped once the error varied by less than 10*^−^*^4^ over five consecutive iterations, after 55 iterations on the canonical microcircuit instance, and the lowest-error matrix, from iteration 51, was applied. Targets above a connection type’s ceiling (Section 1.3) were capped at the emitted ceiling during fitting. Errors reported for the applied network are against the uncapped targets. The filling fractions were fitted on one growth run and applied to a second growth run from the same seed, on which the reported connectivity and its fit are measured. Simulation stochasticity makes the two realisations differ slightly (70,275 against 70,328 neurons), so the fitted fractions transfer to a fresh substrate.

### 3.10 Feasibility analysis and the connection-probability metric

A feasibility pass precedes optimisation. Every f*_ij_* is set to unity and axon–dendrite contacts are counted on the completed substrate. No filling fraction can exceed unity, so the resulting connection probability is the maximum a connection type can reach at any filling fraction. It is a property of the grown morphology alone rather than of the calibration. Pathways whose target exceeds this ceiling are flagged *a priori* as structurally limited, and their residual error after optimisation is a statement about the substrate rather than about the optimiser.

We define the ceiling as a connection probability, that is as the probability that a given source neuron and target neuron share at least one contact. This is the value the equation above takes at f*_ij_* = 1, when every contact becomes a synapse. The quantity the simulation emits is different. It divides the total contact count by the number of possible neuron pairs, so it is a mean contacts-per-pair and exceeds one wherever a few pairs carry many contacts each. Ceilings computed from the emitted quantity therefore understate how many connection types are structurally infeasible. Both quantities count geometric contacts on the completed substrate, not the synapses of the reported network, which are capped at one per connected pair. On the visual instance 5 connection types appear structurally constrained under the emitted metric, against 13 of 64 under the corrected one. We report the corrected metric throughout.

### 3.11 Laminar position tuning

Laminar positions are tuned to match a supplied set of layer depth targets, taken per layer from the rat barrel cortex measurements of Meyer et al. [27], chosen because they are the most complete quantitative laminar measurements available and not as a claim about area identity. Both instances use the same targets; they are an input to the model and not a property of the growth rules. The visual instance differs in its per-layer neuron fractions, set to the composition of the Allen mouse visual cortex model [31]. It was grown before the final adjustment to axonal and layer-4 basal branching (Supplementary Tables S2 and S3) and uses the earlier values of those four parameters, a developmental duration of 10,000 rather than 15,000 steps, and per-iteration sampled rather than closed-form connectivity during fitting. The morphology reference is a third input that could carry area identity, and we hold it generic for both instances.

The migration parameter controls radial displacement, that is vertical position within the column, with higher values producing longer migration and more superficial final positions. It stands in for the local signals that stop migrating neurons at their laminar position during corticogenesis [20]. The resulting parameterisation reproduces the inside-out pattern of cortical development. Early-born neurons destined for L6 carry lower values and settle deeper, while late-born neurons destined for L2/3 carry higher values and settle superficially. Inverting the ordering of these parameters is the developmental lesion used to regrow a reeler-like column.

Migration parameters were adjusted manually, in fewer than ten iterations, by comparing simulated median layer midpoints to the target boundaries. Unlike the automated fit of the connectivity, this is a small manual adjustment of the per-type migration distances. The acceptance tolerance is 100 µm, chosen from the inter-animal variability seen in anatomical studies of rodent cortex. It is smaller than typical layer thicknesses, which run from 150 µm to 600 µm. The tolerance therefore retains biologically meaningful precision while allowing for natural variability.

### 3.12 Spiking network simulation

Spiking simulations used the NEST simulator [32], version 3.9. All neurons used a leaky integrate-and-fire neuron with exponential postsynaptic currents, with membrane parameters taken unchanged from the Potjans and Diesmann reference implementation [7]. Mean excitatory postsynaptic potential amplitude was 0.15 mV and the inhibitory-to-excitatory ratio was g = *−*4, giving a mean inhibitory postsynaptic potential of −0.6 mV. Weight relative standard deviation was 10%, and the postsynaptic current time constant was τ*_syn_* = 0.5 ms. Reference-network delays were drawn per population from Gaussians of mean 1.5 ms for excitatory and 0.75 ms for inhibitory connections, with relative standard deviation 50%.

In the grown network, delays were instead derived per synapse from the morphology. Path distance from the synapse to the presynaptic soma, counted in neurite segments, was divided by a nominal factor of 150 per millisecond, floored at 0.75 ms, then rescaled globally so the mean matched the reference-network value of 1.125 ms. Only the heterogeneity of the delays is morphology-derived, and because the mean is matched by construction the model makes no absolute conduction-delay claim. This uses the geometry the grown network provides and the reference lacks, while matching the mean keeps the two networks’ delays comparable. Distance-proportional delays follow from the finite conduction velocity of local cortical axons, measured directly in the local axons of fast-spiking basket cells [42]. We treat that velocity as uniform across axons, a simplification, since those measurements show it scales with axonal myelination. We retain the heterogeneity deliberately, because spatially uniform delays destabilise the asynchronous irregular state and inflate firing rates. External drive was Poisson input at 8.0 Hz per external synapse, with population-specific external in-degrees. Simulation duration was 1,000 ms, after a discarded 500 ms warm-up, at a resolution of 0.1 ms.

In-degrees and background rate are directly from the canonical Potjans and Diesmann values. Under the weight scheme used here they are scaled by K_ext_ = 0.76 alongside the recurrent connectivity, so the applied external input is that fraction of the canonical level. The scaling is applied identically to the grown and to the size-matched reference network and was not tuned to the grown firing rates. The comparison therefore concerns the difference between the two networks under identical drive, not the absolute rate level of either.

Synaptic polarity is fixed in both networks. During development GABA switches from depolarising in immature neocortex [43, 44] to hyperpolarising as neurons mature [45], a switch that any activity-dependent refinement of the grown wiring would need to include. GABA polarity does not affect the present connectivity, which is set by contacts and filling fractions alone.

### 3.13 Matched scaling procedure

Following an established downscaling procedure [46], we scaled the reference network to the grown one, so that the comparison isolates connectivity structure. Neuron counts are scaled by the ratio of grown to full-scale neurons (70,328 against 77,169, 91%). In-degrees are scaled by the ratio of grown to reference synapses per neuron, which preserves mean in-degree per population (75%).

We apply both factors to the reference network, so the two simulated networks carry identical neuron counts and comparable mean in-degrees. The remaining difference is the recurrent connectivity, which is random in the reference network and grown then calibrated in the grown network. One adjustment is applied to the grown network only: each connection type’s mean weight is multiplied by its target-to-achieved connection-probability ratio (clipped to 0.5–4, computed per column), so that each type delivers the same expected input as in the reference. The comparison therefore tests how the grown wiring is arranged rather than residual calibration error in its strength. Because the reference network is scaled to the same total synapse count, the single-synapse-per-pair representation of the grown network is not expected to bias the comparison.

### 3.14 Firing-rate and irregularity analysis

We computed spiking statistics over a 1,000 ms window, from n = 3 independent NEST realisations of the one fixed grown connectivity. Population mean rates for both networks are given in Fig. 5c. From these we claim only that layer 5 is the most active excitatory population and that inhibition exceeds excitation in every layer, not agreement in absolute rate. Both networks operate above published values under the identically applied external-drive scaling. Mean activity and correlation structure can in principle be preserved under downscaling by an appropriate weight scaling, but only over a limited range of in-degree [46]; we therefore claim no absolute rate level.

We quantified irregularity by the coefficient of variation of the interspike interval, 0.56 to 0.68 in the grown network against 0.58 to 0.67 in the reference network. We quantified synchrony from spike counts in 3 ms bins. Synchrony values at the reduced in-degree simulated here are regime-dependent rather than quantitative predictions.

We computed in-degree distributions directly from the two adjacency matrices. They are descriptive statistics of the wiring, reported as a measured property of the grown substrate. We do not claim them as the demonstrated cause of the rate difference. Establishing that would require resampling the grown connectivity to a reference-like in-degree distribution while holding all else fixed, which we have not done.

### 3.15 Perturbation protocols

Synaptic perturbations thin the realised synapses of the affected connection types on the same unchanged wiring, keeping each with probability s (fixed thinning seed), with no regrowth and no retuning of neuron or synapse parameters. We applied two motifs, an excitatory-to-inhibitory knockdown [34, 35] and an inhibitory-to-excitatory knockdown [5], each at *×*0.7, *×*0.5 and *×*0.3 of baseline, and averaged over 3 seeds. Both are restricted to the four within-layer connection types of the relevant class rather than to all cross-layer pairs, because a global cross-layer lesion saturates the network instead of grading it. The sweep’s baseline is the same n = 3 healthy run as Fig. 5 (layer-5 excitatory 15.55 Hz). Rates were computed from the neurons whose spike records were intact (80–97% per run).

The developmental-axis perturbation instead inverts the migration-parameter ordering and regrows the column from progenitors under otherwise identical rules. Connectivity is then recomputed and simulated by the same pipeline. As a control, the size-matched reference network is rescaled to the raised synapse count of the lesioned network, which separates the contribution of connectivity structure from that of synapse count alone.

### 3.16 Use of generative AI

Generative AI tools (Claude, Anthropic) were used to assist with drafting and editing text and with writing analysis and simulation code. All AI-assisted text and code were reviewed, tested and verified by the authors, who take full responsibility for the content of this manuscript.

## Acknowledgements

We thank Rodney Douglas for helpful comments and discussion, Christoph von der Malsburg and Marco Manca for helpful feedback on the manuscript, and Akram Dweikat for donated computing resources. We are grateful to the BioDynaMo consortium for their help with the simulation platform and for feedback throughout this work.

## Data availability

The standardised SWC morphologies, optimised connectivity matrices, per-iteration optimisation metrics and spiking outputs supporting this study will be deposited in a public repository with a citable DOI on publication. The deposit includes the parameter configurations and random seeds needed to reproduce them. The data underlying every figure are provided to editors and reviewers with the code during peer review. Reference data are drawn from NeuroMorpho.org [28], the Potjans and Diesmann supplementary materials [7] and the Allen Institute mouse visual cortex model [31]. Supplementary Videos 1 and 2 are available at https://doi.org/10.5281/zenodo.23675065.

All structural results, meaning morphology, connectivity and the 205-million-pair network, derive from a single grown realisation of the canonical column, apart from the contact counts and feasibility ceilings, which come from the fitting run (Methods). The n = 3 spiking ensembles are independent NEST simulations of that one fixed connectivity rather than independent regrowths. Because growth is stochastic, independent realisations differ in fine detail while sharing the same population-level statistics by construction. We report one realisation and do not quantify across-realisation variability here.

## Code availability

The developmental growth simulation uses BioDynaMo [22] (https://biodynamo.org), built from fork commit bd0671d (v1.05) by the Dockerfile supplied with the code. Spiking simulations use the NEST simulator [32]. The growth simulation, the filling-fraction optimiser and the validation and analysis pipeline are available to editors and reviewers during peer review, and will be released as an open-source repository with a citable DOI on publication, under the Apache-2.0 licence (scripts derived from the NEST microcircuit example: GPL-2.0-or-later). That release will enable adaptation to other species, cortical areas and developmental conditions.

## Author contributions

U.A. wrote all simulation and analysis code, performed the simulations and analyses, and wrote the manuscript. C.H. and R.Bo. contributed to the development of ideas and edited sections of the manuscript. R.Ba. supervised the project and edited the manuscript. All authors reviewed and approved the final manuscript.

## Supplementary information

Supplementary Fig. S1 and Tables S1–S3 are provided as a separate document.

