## Supplementary Information for "Computational model of developmental self-assembly of a cortical microcircuit"

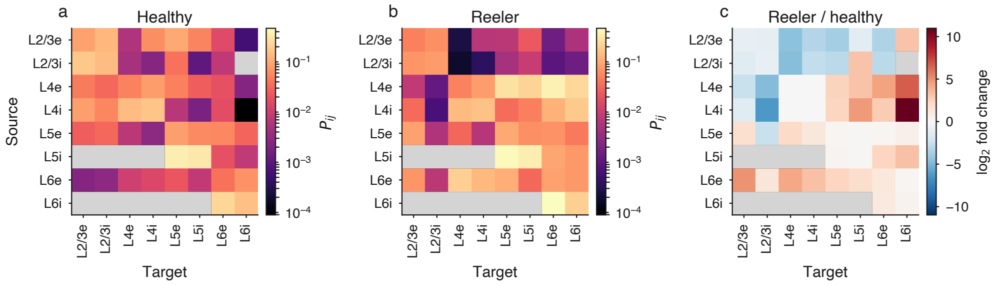


**Fig. S1 Mislamination generates aberrant cross-layer excitatory connectivity.** Connection probability between the eight cortical populations (source × target) for (a) the canonical microcircuit instance and (b) a reeler-like column regrown under inverted lamination and otherwise identical growth and synapse-formation rules, both on a shared logarithmic scale. Grey cells carry no connection; in (c) grey also marks a ratio that is undefined because the canonical column lacks that connection. (c) $\text{Log}_2$ ratio of the two, red where the lesioned column connects more strongly. Mean connection probability rises from 0.049 to 0.078, and the increases concentrate in projections from layer-4 and layer-6 sources, especially onto layer 6, whereas supragranular sources connect less. No single-neuron or synaptic parameter was retuned between the two.

**Table S1 Laminar organisation of the canonical microcircuit instance.** Left: achieved median soma depth per layer against the depth targets used to tune migration, taken from rat barrel cortex [1] (layer boundaries 0, 75, 246, 518, 781, 1289 and 1774 µm). Targets are inputs, not properties of the growth rules. Right: neuron density per layer, as a percentage of L2–L6 neurons, for the Potjans and Diesmann targets [2] and the grown column; per-layer apoptosis rates bring the grown fractions to these targets. The Potjans L2/3 value (34.4%) is shared by the L2 and L3 rows (grown L2+L3 = 34.2%). MZ/L1 has no network population, hence the dashes.

| | Layer position (µm) | | | Neuron density (%) | |
|---|---|---|---|---|---|
| Layer | Target | Achieved | Error | Potjans | Grown |
| MZ/L1 | 40 | ∼42 | 2 | — | — |
| L2 | 200 | ∼208 | 8 | 34.4 | 18.0 |
| L3 | 437 | ∼380 | 57 | | 16.1 |
| L4 | 525 | ∼536 | 11 | 35.5 | 36.0 |
| L5 | 1006 | ∼978 | 28 | 7.7 | 7.3 |
| L6 | 1548 | ∼1395 | 153 | 22.5 | 22.5 |

**Table S2 Growth-cone parameters, excitatory neurons.** The gradient weight scales how strongly the cone turns up the guidance substance; the random weight scales the stochastic component of each step. Bifurcation probability is evaluated per growth step. Diameter loss is the per-step taper, which terminates a branch when its diameter reaches the minimum. Values are read directly from the simulation configuration used for the reported run.

| Compartment | Layer | Gradient weight | Random weight | Bifurcation probability | Diameter loss (per step) |
|---|---|---|---|---|---|
| Axon | MZ | 2 | 1.5 | 0 | $4.7 \times 10^{-4}$ |
| | L2 | 1.5 | 0.8 | $1.38 \times 10^{-4}$ | $7.16 \times 10^{-5}$ |
| | L3 | 1.5 | 0.8 | $3.75 \times 10^{-4}$ | $8 \times 10^{-5}$ |
| | L4 | 1.2 | 0.8 | $3.8 \times 10^{-5}$ | $8.68 \times 10^{-5}$ |
| | L5 | 1.5 | 0.8 | $8.5 \times 10^{-5}$ | $1 \times 10^{-4}$ |
| | L6 | 2 | 0.8 | $2.2 \times 10^{-5}$ | $8 \times 10^{-5}$ |
| Apical | MZ | 0.3 | 0.3 | $9.38 \times 10^{-5}$ | $8 \times 10^{-6}$ |
| | L2 | 0.3 | 0.3 | $3 \times 10^{-5}$ | $2.2 \times 10^{-4}$ |
| | L3 | 0.3 | 0.3 | $3.5 \times 10^{-5}$ | $1.25 \times 10^{-4}$ |
| | L4 | 0.3 | 0.3 | $6.5 \times 10^{-5}$ | $1.7 \times 10^{-4}$ |
| | L5 | 0.3 | 0.3 | 0.001 | $1.6 \times 10^{-4}$ |
| | L6 | 0.3 | 0.3 | 0.002 | $4.5 \times 10^{-5}$ |
| Basal | MZ | 0.03 | 1.2 | 0.0026 | $4.6 \times 10^{-4}$ |
| | L2 | 0.03 | 2 | 0.003 | $6 \times 10^{-4}$ |
| | L3 | 0.03 | 2 | 0.0035 | $6 \times 10^{-4}$ |
| | L4 | 0.03 | 2 | 0.003 | $8 \times 10^{-4}$ |
| | L5 | 0.03 | 2 | 0.004 | $8 \times 10^{-4}$ |
| | L6 | 0.03 | 2 | 0.0045 | $8 \times 10^{-4}$ |

**Table S3 Growth-cone parameters, inhibitory neurons.** Columns as in Table S2. Values are for basket cells, except the apical rows, which apply to Martinotti cells, the only interneurons that grow an apical dendrite. Martinotti axons use their own direction weights and apply fixed scale factors to the axonal bifurcation and taper values shown.

| Compartment | Layer | Gradient weight | Random weight | Bifurcation probability | Diameter loss (per step) |
|---|---|---|---|---|---|
| Axon | MZ | 0.3 | 1.5 | $4 \times 10^{-6}$ | 0.0012 |
| | L2 | 0.4 | 1.5 | $2.25 \times 10^{-4}$ | $1.34 \times 10^{-4}$ |
| | L3 | 0.4 | 1.5 | $2.1 \times 10^{-4}$ | $1.25 \times 10^{-4}$ |
| | L4 | 0.3 | 1.5 | $1.65 \times 10^{-4}$ | $1.05 \times 10^{-4}$ |
| | L5 | 0.2 | 1.5 | $1.3 \times 10^{-4}$ | $1.1 \times 10^{-4}$ |
| | L6 | 3 | 0.4 | $1.5 \times 10^{-4}$ | $1.6 \times 10^{-4}$ |
| Apical | MZ | 0.3 | 2 | 0 | $3 \times 10^{-4}$ |
| | L2 | 0.3 | 2 | $5 \times 10^{-5}$ | $3 \times 10^{-4}$ |
| | L3 | 0.3 | 2 | $6 \times 10^{-5}$ | $2 \times 10^{-4}$ |
| | L4 | 0.3 | 2 | $3 \times 10^{-4}$ | $4.5 \times 10^{-4}$ |
| | L5 | 0.3 | 2 | 0.002 | $1.5 \times 10^{-4}$ |
| | L6 | 0.9 | 2 | 0.0035 | $2.2 \times 10^{-4}$ |
| Basal | MZ | 0.03 | 0.05 | 0.002 | $4.6 \times 10^{-4}$ |
| | L2 | 0.03 | 0.04 | 0.003 | $8 \times 10^{-4}$ |
| | L3 | 0.03 | 0.04 | 0.0031 | $8 \times 10^{-4}$ |
| | L4 | 0.03 | 0.05 | 0.0035 | 0.001 |
| | L5 | 0.03 | 0.04 | 0.0035 | $8 \times 10^{-4}$ |
| | L6 | 0.03 | 0.02 | 0.0031 | $8.8 \times 10^{-4}$ |

### Supplementary Videos

**Supplementary Video 1. Self-assembly of the cortical column.** Time-lapse of the column self-assembling, from progenitor proliferation and inside-out lamination to neurite growth. Only a small fraction of neurons grow neurites in this visualisation run of the same growth rules at reduced growth fraction, as in Fig. 1c.

**Supplementary Video 2. Growth-cone tracking of an ascending axon.** A camera follows the growth cone of the ascending axon of a layer-5 pyramidal cell as it climbs about 700 μm through the column, from the same visualisation run. Neighbouring cells are shown for context.

Both videos are provided as separate files and are available at https://doi.org/10.5281/zenodo.23675065.
